# Zika Virus Subverts Stress Granules to Promote and Restrict Viral Gene Expression

**DOI:** 10.1101/436865

**Authors:** Gaston Bonenfant, Nina Williams, Rachel Netzband, Megan C. Schwarz, Matthew J. Evans, Cara T. Pager

## Abstract

Flaviviruses limit the cell stress response by preventing the formation of stress granules and modulate viral gene expression by subverting different proteins involved in the stress granule pathway. In this study, we investigated the formation of stress granules during Zika virus (ZIKV) infection and the role stress granule proteins play during the viral life cycle. Using immunofluorescence and confocal microscopy, we determined that ZIKV disrupted the formation of arsenite-induced stress granules and changed the subcellular distribution, but not the abundance or integrity, of stress granule proteins. We also investigated the role of different stress granule proteins in ZIKV infection by using target-specific siRNAs to deplete Ataxin2, G3BP1, HuR, TIA-1, TIAR and YB1. Knock-down of TIA-1 and TIAR affected ZIKV protein and RNA levels, but not viral titers. Conversely, depletion of Ataxin2 and YB1 decreased virion production despite having only a small effect on ZIKV protein expression. Notably, however, depletion of G3BP1 and HuR decreased and increased ZIKV gene expression and virion production, respectively. Using an MR766 *Gaussia* luciferase reporter genome together with knockdown and overexpression assays, G3BP1 and HuR were found to modulate ZIKV replication. These data indicate that ZIKV disrupts the formation of stress granules by sequestering stress granule proteins required for replication, where G3BP1 functions to promote ZIKV infection, while HuR exhibits an antiviral effect. The consequence of ZIKV re-localizing and subverting select stress granule proteins might have broader consequences on cellular RNA homeostasis and contribute to cellular gene dysregulation and ZIKV pathogenesis.

**Importance:** Many viruses inhibit stress granules (SGs). In this study, we observed that ZIKV restricts SG assembly likely by re-localizing and subverting specific SG proteins to modulate ZIKV replication. This ZIKV-SG protein interaction is interesting, as many SG proteins are also known to function in neuronal granules, which are critical in neural development and function. Moreover, dysregulation of different SG proteins in neurons has been shown to play a role in the progression of neurodegenerative diseases. The likely consequences of ZIKV modulating SG assembly and subverting specific SG proteins are alterations to cellular mRNA transcription, splicing, stability, and translation. Such changes in cellular ribostasis could profoundly affect neural development and contribute to the devastating developmental and neurological anomalies observed following intrauterine ZIKV infection. Our study provides new insights into virus-host interactions and the identification of the SG proteins that may contribute to the unusual pathogenesis associated with this re-emerging arbovirus.

## Introduction

Zika virus (ZIKV) is an enveloped, single-stranded positive-sense RNA virus belonging to the *Flaviviridae* family, which includes Dengue virus (DENV), Yellow fever virus (YFV), and West Nile virus (WNV) (1). While ZIKV was discovered in Uganda in 1947 (2), the virus garnered renewed interest during the 2015-2016 outbreak in the Americas (3), in particular because of intrauterine infections and resulting developmental abnormalities such as severe microcephaly, decreased brain tissue, macular scarring, congenital contractures, and hypertonia (4–9). Additionally, adults infected with ZIKV were reported to develop Guillain-Barré syndrome, a debilitating disorder affecting the peripheral nerves (10–13). Similar to other flaviviruses, ZIKV is transmitted by the *Aedes aegypti* and *Aedes albopictus* mosquitoes, although recent evidence has shown ZIKV is also capable of sexual and vertical transmission (14–17). While half a century has passed since the discovery of ZIKV, little-to-no research was published prior to the emergence of the current strain in the Americas associated with devastating developmental pathologies. Because there is no licensed vaccine and antiviral treatments are elusive, a fundamental understanding of the molecular biology of ZIKV and virus-host interactions is critical to developing therapeutic strategies.

The ZIKV single-stranded positive-sense RNA genome contains a 5’-cap, lacks a poly(A) tail, and encodes one open reading frame (ORF) that is flanked by highly structured 5’ and 3’ untranslated regions (UTRs). Similar to other flaviviruses, translation of the ZIKV RNA results in one long polyprotein that is co- and post-translationally proteolytically processed to produce at least three structural proteins (capsid [C], premembrane [prM] and envelope [E]) and seven nonstructural proteins (NS1, NS2a, NS2b, NS3, NS4a, NS4b and NS5) (1). Although cap-dependent and cap-independent translation have been reported for DENV (18), it is presently unknown whether ZIKV employs similar translation strategies. Similarly, little is known regarding the strategies ZIKV employs to promote translation of the viral RNA.

To limit translation of viral RNAs, or protect the cells from different environmental stresses, mammalian cells rapidly stall translation via the activation of one of the four eIF2α kinases. In particular, the presence of double-stranded RNA (dsRNA) during viral infection activates protein kinase R (PKR) (19); the accumulation of unfolded proteins in the endoplasmic reticulum (ER) and resulting stress activates PKR-like endoplasmic reticulum kinase (PERK) (20); amino acid starvation activates general control non-repressed 2 (GCN2) (21); and oxidative stress activates heme-regulated inhibitor kinase (HRI) (22). Phosphorylation of the α subunit of eIF2 by one of the four stress response kinases results in the stalling of translation initiation, and disassembly of polysomes. Stalled translation initiation complexes bound to mRNA are recognized by several RNA binding proteins, which aggregate to form RNA-protein macromolecular complexes called stress granules (SGs) (23). Once the stressor is abated, eIF2α is dephosphorylated by protein phosphatase 1 (PPI) and the PPI cofactor growth arrest and DNA-damage-inducible 34 (GADD34), allowing for the return of sequestered mRNA transcripts to active translation (23).

SGs are dynamic nonmembrane-bound cytoplasmic structures that can rapidly assemble in response to stress and disassemble once the stress has been alleviated (23). SGs typically contain mRNAs, stalled translation initiation complexes, and numerous RNA binding proteins. Indeed, SGs may contain upwards of 260 different proteins (24), and ~50% of these are proposed to be RNA-binding proteins (25). Of these Ras-GTPase activating binding protein 1 (or GAP SH3 domain binding protein 1 [G3BP1]), Caprin1, T-cell internal antigen 1 (TIA-1) and TIA-1 related protein (TIAR) are proposed to be key nucleators of SG assembly (26–29). In addition to translation repression and mRNA sorting, SGs also amplify the innate immune response by aggregating critical antiviral factors (23). Because translation is a critical first step in the flavivirus life cycle, the formation of SGs presents an immediate obstacle to infection. Notably, however, during infection with different flaviviruses, such as WNV, DENV, and Japanese encephalitis virus (JEV) SGs are absent, and treatment of virus-infected cells with arsenite fails to induce SGs (30–33). While WNV, DENV and JEV all belong to the same flavivirus genus, each virus employs a unique mechanism to block SG assembly. For example, early during WNV infection PKR is activated by the appearance of exposed dsRNA replication intermediates, which results in phosphorylation of eIF2α, the stalling of translation initiation and formation of stress granules. However, at later times during infection SG formation is limited as membranous vesicles mask dsRNA and viral replication complexes, and thus PKR remains inactive (30, 34, 35). More recently, WNV was shown to also limit SG assembly by upregulating and activating key transcription factors that modulate the antioxidant response (32). In contrast, p38-Mnk1 signaling and phosphorylation of the cap-binding protein eIF4E was affected during DENV infection, thus inhibiting SG formation via an eIF2α phosphorylation-independent mechanism (33). Flaviviruses also subvert specific SG proteins to promote viral gene expression. For example, the SG-nucleating proteins TIA-1 and TIAR facilitate WNV replication (36); G3BP1, G3BP2 and Caprin1 promote translation of interferon stimulated genes (ISGs) to limit DENV infection (37, 38); and JEV inhibits SG assembly by co-localizing Caprin1 with the viral capsid protein (31).

Similar to DENV and WNV, ZIKV was recently shown limit the assembly of SGs (39, 40). In particular, Hou *et al.* reported that exogenously expressed Flag-tagged NS3 and NS4A repressed translation, and the formation of SGs was restricted when Flag-tagged capsid, NS3, NS2B-3 and NS4A proteins were individually expressed (39). Moreover ZIKV RNA was bound by G3BP1, and G3BP1 and Caprin1 co-immunoprecipitated Flag-tagged ZIKV capsid protein (39). Depletion of TIAR, G3BP1 and Caprin1 decreased viral titers, likely as a result of disrupting the interactions with ZIKV capsid protein and RNA (39). Hou *et al.*, also showed that SG assembly in ZIKV-infected cells was the consequence of eIF2α phosphorylation and translational repression by activating PKR and the unfolded protein response (39). Amorim et al. additionally reported that ZIKV limits the antiviral stress response by promoting an increase in the rate of eIF2α dephosphorylation during infection (40). While these studies provide insight into how SG formation is inhibited during ZIKV infection, the role of various SG proteins on ZIKV gene expression has yet to be elucidated.

A number of the RNA-binding proteins that localize in SGs, such as Ataxin-2, G3BP1, HuR and TIA-1, are known to contribute to different neuropathologies (41, 42). Thus, our goal in this study was to investigate the biological impact of different SG proteins linked to neurodegeneration on ZIKV gene expression. Here we show a systematic analysis of SGs during ZIKV infection; the effect of depleting six different SG proteins on ZIKV protein and RNA levels and viral titers; and elucidate the biological function of two SG proteins on ZIKV gene expression. We determined that ZIKV disrupted arsenite-induced SG assembly and that several SG markers co-localized with sites of ZIKV replication. Additionally, the SG protein G3BP1 is required for ZIKV gene expression, while HuR exhibited antiviral activity. Using a MR766 *Gaussia* luciferase reporter genome, our studies revealed that G3BP1 and HuR specifically modulate ZIKV replication. This work advances our understanding of the interplay between ZIKV, the cellular stress response and cellular RNA metabolism, and demonstrates a role for specific RNA-binding proteins in ZIKV replication.

## Materials and Methods

### Cell Maintenance and ZIKV Stocks

The hepatocellular carcinoma cell line Huh7 was maintained in Dulbecco’s minimal essential medium (DMEM; Life Technologies) supplemented with 10% fetal bovine serum (FBS; Seradigm), 10 mM nonessential amino acids (NEAA; Life Technologies), and 5 mM L-glutamine (Life Technologies). Vero cells (ATCC# CRL–81) were maintained in DMEM supplemented with 10% FBS and 10 mM HEPES (Life Technologies). Mammalian cell lines were grown at 37°C with 5% CO_2_. C6/36 cells (ATCC# CRL-1660) grown at 27°C with 5% CO_2_ were maintained in Eagle’s minimal essential medium (EMEM; Sigma) supplemented in 10% FBS, sodium pyruvate (0.055 g/L; Life Technologies), Fungizone (125 μg/L; Life Technologies), and penicillin and streptomycin (50000 units/L penicillin, 0.05 g/L streptomycin; Life Technologies). ZIKV (Cambodia-160310, Uganda MR766, and Puerto Rico PRVABC59; generous gifts from Dr. Brett Lindenbach, Yale School of Medicine; Dr. Laura Kramer, Wadsworth Center NYDOH; and the CDC) stocks were generated in C6/36 cells. Briefly, C6/36 cells nearing confluency were infected at a multiplicity of infection (MOI) of 0.1. Seven days post-infection supernatants from the infected cells were collected, aliquoted and stored at −80°C. RNA was extracted from cells to confirm infection via northern blotting, and viral titers were determined by plaque assay.

### Small Interfering RNA (siRNA) and DNA Plasmid Transfections

Sense and antisense siRNA oligonucleotides were synthesized by Integrated DNA Technologies (IDT). The siRNA sequences are provided in Table 1. Oligonucleotides were resuspended in RNase-free water to a 50 μM final concentration. Sense and antisense strands were combined in annealing buffer (150 mM HEPES (pH 7.4), 500 mM potassium acetate, and 10 mM magnesium acetate) to a final concentration of 20 μM, denatured for 1 minute at 95°C, and annealed for 1 hour at 37°C (43).

**Table 1:**
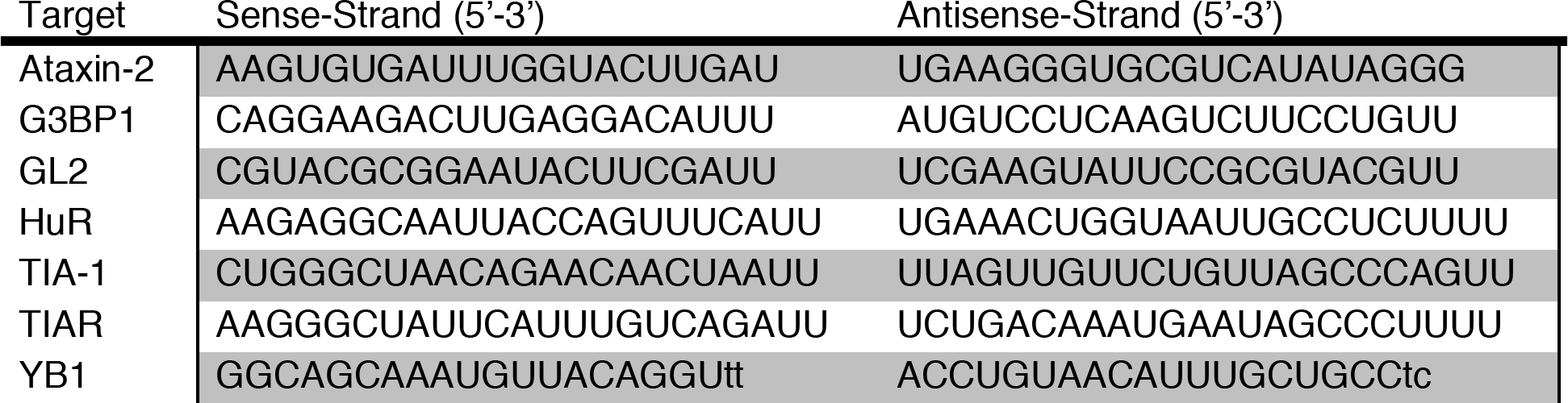
siRNA Sequences

For siRNA and plasmid transfections, Huh7 cells seeded at 7.5×10^6^ in a 10 cm cell culture dish were transfected 24 hours later with 100 nM of the indicated siRNA duplex or 2 μg of 3xFLAG-BAP (Bacterial Alkaline Phosphatase; Sigma #C7472), HuR-Flag (GenScript #OHu23723D), or G3BP1-Flag (GenScript #OHu02150D) plasmids using Lipofectamine 3000 (Invitrogen) per manufacturer’s protocol. At 24 hours post-transfection, cells were infected with ZIKV.

### ZIKV Infections

On the day of infection, one mock culture plate was trypsinized and counted to calculate MOI. Cells were infected at either MOI of 1 or 5, as noted in the text and figure legends. Appropriate amount of viral stocks thawed at room temperature were diluted in PBS to a final volume of 3.5 mL. Culture dishes were aspirated, the viral solution was added, and cells were returned to the incubator for 1 hour with rocking every 15 minutes. At the end of the hour 4 mL complete media was added to each plate.

### Immunofluorescence Analysis and Confocal Microscopy

Cells seeded in 8-chambered slides (Nunc Lab-Tek Chamber Slide system; Sigma C7182) were infected with ZIKV at MOI of 5. Two days post-infection, cells were either mock-treated or treated with sodium arsenite (1 mM final concentration; Sigma) for 30 minutes. Hereafter the cells were washed three times with phosphate-buffered saline (PBS) and fixed with paraformaldehyde (4%)-PBS for 10 minutes at room temperature. Cells were washed once with PBS and permeabilized with 100% ice-cold methanol for 15 minutes at room temperature. Cells were washed in blocking buffer (PBS-1% fish gelatin (FG); Sigma) three times for 15 minutes. Primary antibody was diluted in blocking buffer, added to the appropriate wells and incubated overnight at 4°C. Secondary antibodies diluted in blocking buffer were added for 1 hour at room temperature in the dark. Hoechst-33342 (Life Technologies) was applied for 15 minutes. Between and after application of antibodies, cells were washed with blocking buffer three times for 15 minutes. Finally, the cells were washed twice with PBS for 5 minutes, the 8-chamber upper structure was removed, Fluoromount (Southern Biotech) added, and a cover-slip applied. Slides were stored at 4°C if imaging was undertaken within 1 week, or at −20°C for long-term storage. Antibodies and concentrations used are listed in Table 2. Slides were imaged on a Zeiss LM710 confocal microscope with a 63x oil objective.

**Table 2:** Antibodies and Concentrations

| Protein | Supplier (Product ID) | Species | IFA Dilution | Western Dilution |
| --- | --- | --- | --- | --- |
| G3BP1 | Bethyl (A302-034A) | Rabbit | 1:300 | 1:5,000 |
| TIAR | Bethyl (A303-613A) | Rabbit | 1:250 | 1:1,000 |
| YB1 | Cell Signaling (D299) | Rabbit | 1:250 | N/A |
| Ataxin-2 | ProteinTech (21776-1-AP) | Rabbit | 1:200 | 1:5,000 |
| HuR | ProteinTech (11910-1-AP) | Rabbit | 1:200 | 1:6,000 |
| YB1 | Novus (NBP1-97572) | Rabbit | N/A | 1:5,000 |
| TIA-1 (C-20) | Santa Cruz (Sc-1751) | Goat | 1:50 | 1:3,000 |
| eIF3 | Santa Cruz (Sc-16377) | Goat | 1:50 | N/A |
| GRP78 | Santa Cruz (Sc-376768) | Goat | N/A | 1:1,000 |
| Fibrillarin | ProteinTech(16021-1-AP) | Rabbit | N/A | 1:5,000 |
| Calnexin | ProteinTech (10427-1-AP) | Rabbit | N/A | 1:5,000 |
| GAPDH | Millipore Sigma | Mouse | N/A | 1:10,000 |
| ZIKV Capsid | GeneTex (GTX133317) | Rabbit | N/A | 1:5,000 |
| ZIKV Envelope | BioFront (1176-76) | Mouse | 1:800 | N/A |
| ZIKV NS5 | BioFront (6A1) | Mouse | 1:150 | N/A |
| Anti-rabbit-<br>peroxidase | Santa Cruz (Sc-2313) | Donkey | N/A | 1:10,000 |
| Anti-goat-<br>peroxidase | Santa Cruz (Sc-2354) | Mouse | N/A | 1:5,000 |
| Anti-mouse-<br>peroxidase | Santa Cruz (Sc-2314) | Donkey | N/A | 1:10,000 |
| Anti-mouse-488 | Life Technologies | Donkey | N/A | 1:200 |
| Anti-rabbit-594 | Life Technologies | Donkey | N/A | 1:200 |
| Anti-goat-647 | Life Technologies | Donkey | N/A | 1:200 |

\*N/A Not applicable

### Quantification of SG-positive Cells

A 40x oil objective was used to acquire images of ZIKV-infected cells stained with mouse-anti-dsRNA antibody to detect ZIKV-infected cells and one of the following anti-SG protein antibodies: rabbit-anti-Ataxin-2, rabbit-anti-G3BP1, rabbit-anti-HuR, goat-anti-TIA-1, rabbit-anti-TIAR, and rabbit-anti-YB1. To quantify the effect of ZIKV infection on SG formation, 200 to 300 infected cells between three or more biological replicates were counted. Quantification of SGs between different infections was completed as follows: for each treatment, the percentage of uninfected cells with/without SGs and ZIKV-infected cells with/without SGs was calculated. Using the percentages from cells with/without SGs from three or more biological replicates, a one-tailed student T-test was carried out.

### Subcellular Fractionation

Huh7 cells seeded in 10 cm cell culture dishes at 7.5×10^6^ were infected at a MOI of 5. Two days post-infection, the cells were trypsinized, centrifuged at 500 x g for 5 minutes, and resuspended in PBS. Cytoplasmic and nuclear proteins were separated using the NE-PER extraction reagents per manufacturer’s protocol (ThermoScientific). After clarification of proteins, 30 μg of protein lysate from the cytoplasmic and nuclear fractions were analyzed by western blot analysis. To demonstrate successful separation of cytoplasmic and nuclear fractions the western blots were also probed for the ER chaperone protein GRP78 and Fibrillarin, a methyltransferase enzyme that 2’O-methylates ribosomal RNA, respectively. Quantification of band intensities was performed using Image Lab software (BioRad). The subcellular distribution of SG proteins in ZIKV-infected cells was normalized to the corresponding mock-infected lane, and an average of three independent experiments was used to generate the quantification graph.

### Harvest of ZIKV-Infected Cells for Protein and RNA Analysis

At the indicated time point, media from culture dishes was aspirated and the cells were washed once with PBS. Cells were manually scraped from plate in 1 mL PBS, divided between two microcentrifuge tubes, and pelleted at 14,000 RPM for 15 seconds. The supernatant was removed, and cells were resuspended in 50 μL RIPA buffer (100 mM Tris-HCl (pH 7.4), 150 mM NaCl, 1% sodium deoxycholic acid, 1% Triton-X100, 0.1% SDS) containing protease inhibitors (mini tablets EDTA-free; ThermoScientific Pierce) or 1 mL TRIzol (Invitrogen) for protein and RNA analysis, respectively. For protein analysis, cells resuspended in RIPA buffer with protease inhibitors were incubated on ice for 20 minutes. The lysate was clarified by centrifugation at 14,000 RPM at 4°C for 20 minutes. The clarified protein lysate was collected and processed for western blot analysis. RNA from cell lysates was extracted using TRIzol per the manufacturer’s instructions.

### Western Blot Analysis

Protein lysates were quantified using the DC protein assay kit (BioRad) and 20 μg lysate was electrophoresed for 2 hours at 100V through SDS-10% PAGE. Protein was transferred to a PVDF membrane at 100 V for 60 minutes at 4°C. Transfer efficiency was determined by Ponceau S (Sigma) staining, after which the blots were washed in buffer (0.1% Tween in PBS (PBS-T)) and blocked for 30 minutes in blocking buffer (5% powdered milk (w/v) in PBS-T). Primary antibodies diluted in blocking buffer were added to the blots and incubated at 4°C overnight. Secondary antibodies diluted in blocking buffer were incubated at room temperature for 1 hour. The assay was developed using Clarity Western ECL Blotting Substrate (BioRad) and imaged on a chemiluminescent imager (BioRad). Prior to application of additional antibody, blots were stripped using ReBlot Plus Mild (Millipore Sigma) according to manufacturer’s suggestion. Before and after application of primary/secondary antibodies and stripping, blots were washed for 15 minutes three times in wash buffer. Antibodies and concentrations are listed in Table 2.

### Northern Blot Analysis

TRIzol-extracted RNA (10 μg) was resuspended in loading buffer (1x MOPS-EDTA-sodium acetate (MESA; Sigma), 4.5% formaldehyde, and 32% formamide), denatured for 15 minutes at 65°C, separated in a 1.2% agarose gel containing 5.92% formaldehyde (v/v) and 1x MESA (v/v), and transferred via capillary action to a Zeta-probe membrane (BioRad) overnight at room temperature. RNA was crosslinked to the membrane using a TL-2000 (UVP) and then stained with methylene blue to visualize transfer of RNA. Methylene blue was removed by washing the membranes in 1xSSC/1% SDS for 15 minutes three times, and then prehybridized in 5 mL ExpressHyb (ClonTech) for 1 hour at 65°C. The hybridization buffer was changed and a ^32^P-labeled dsDNA probe (Invitrogen) was added for 1 hour at 65°C. Blots were then washed with 0.1xSSC/0.1% SDS three times for 15 minutes at 55°C, exposed overnight to a phosphorimager screen, and subsequently visualized on a Typhoon 9400 (GE). The ZIKV 3’ UTR probe (targeting nt 10324-10808 of the viral genome) was generated by RT-PCR amplification of the 3’ UTR region from ZIKV-infected cells and the resulting PCR product cloned into pCR-TOPO2.1 (Invitrogen). The actin and ZIKV probes were randomly labeled with α^32^P dATP (Perkin Elmer) using a RadPrime labeling kit (Invitrogen).

### Isolation of ZIKV Replication Complexes

A protocol adapted from Schlegel *et al.*, and Chen *et al.*, was used to isolate ZIKV replication complexes (44, 45). Huh7 cells seeded in 15 cm cell culture dishes were either mock-infected or infected with ZIKV (Cambodia-160310) at a MOI of 1 and then harvested 1-day post-infection. Specifically, cells were washed twice with cold PBS, gently dislodged using a cell lifter, collected and pelleted at 1,000 RPM for 5 minutes at 4°C. The cell pellets were resuspended in 1 mL hypotonic buffer (10 mM Tris-HCl [pH 8.0], 10 mM NaCl, 1 mM MgCl_2_) containing protease and phosphatase inhibitors (mini tablets EDTA-free; ThermoScientific Pierce). One-tenth volume of cells (100 μL) was collected for the protein input sample, which was lysed in RIPA buffer as described above. The cells in hypotonic buffer were incubated on ice for 10 minutes. After incubation, cells were homogenized using the tight pestle of a Dounce homogenizer approximately 50 times on ice. Samples were then centrifuged at 1,000 x g for 10 minutes at 4°C, the supernatant collected and NaCl was added to the supernatants to a final concentration of 300 mM. The replication complexes in the collected supernatant were separated in a 10%/60% gradient. The 10%/60% sucrose gradient was prepared in Ultraclear tubes (Beckman Coulter) by first adding 5.5 mL of 10% sucrose solution (300 mM NaCl, 15 mM Tris-HCl [pH 7.5], 15 mM MgCl_2_, and 10% sucrose), followed by a slow deposition of 5.5 mL 60% sucrose (300 mM NaCl, 15 mM Tris-HCl [pH 7.5], 15 mM MgCl_2_, and 60% sucrose) below the 10% sucrose layer. Samples were layered onto the sucrose gradients and ultracentrifuged using the SW41 rotor at 26,000 RPM for 16 hours at 4°C. The opaque layer sedimenting between the 10% and 60% sucrose layers was collected for analysis. For western blot analysis the proteins were precipitated with methanol and resuspended in a buffer containing 8 M urea and 100 mM Tris-HCl (pH 8.0).

### Plaque Assays

Six-well plates were seeded with 6×10^5^ Vero cells per well. The following day, serial dilutions of virus were prepared in PBS, media was aspirated, and 300 μL of appropriate dilution was added. Cultures were returned to incubator for 1 hour with rocking every 15 min. After incubation, 2.5 mL of a 1:1 overlay (1.2% Oxoid agar and modified media (2xDMEM, 4% FBS, 10 mM HEPES) was added to each well. Agar was left to solidify at room temperature for 10 minutes before returning the plates to incubator. At 4 days post-infection, plaques were developed using 1% crystal violet in 20% methanol.

### Cell Viability Assays

Huh7 cells seeded at 5×10^5^ in a 6 cm cell culture dish were transfected 24 hours later with the indicated siRNA duplex using Lipofectamine 3000 (Invitrogen) as per manufacturer’s protocol. After 24 hours, cells were trypsinized and seeded in triplicate into a white 96-well plate. Forty-eight hours after seeding the multi-well plate, the original 6 cm culture dish was harvested for protein to confirm knockdown, and a cell viability assay was performed. An equal volume of CellTiter-Glo 2.0 (100 μL; Promega) to cell culture media was added to each well, which was rocked for two minutes following a ten-minute room temperature incubation. Luminescence was recorded with an integration time of one second using a BioTEK Synergy luminometer.

### Luciferase Reporter Virus and Assays

To construct a ZIKV reporter virus, the *Gaussia* luciferase gene was cloned as a fusion with the amino terminus of the polyprotein, as previously described for another ZIKV strain (46), into the previously described plasmid pCDNA6.2 MR766 Intron3127 HDVr encoding a CMV promoter driven MR766 ZIKV (47). Specifically, a translational fusion was generated comprised of the first 20 amino acids of the Capsid protein, the full-length *Gaussia* luciferase reporter gene, along with its signal sequence, the FMDV 2A peptide, and finally the full-length viral polyprotein. The flavivirus cyclization sequence determinants in full length capsid coding region was disrupted by silent mutagenesis (indicated here by lower case nucleotides: 5’ ATT GTa AAc ATG tTA AAA) To create a replication incompetent version of this plasmid, this reporter cassette was also cloned into the previously described pCDNA6.2 MR766 Intron3127 Pol(-) HDVr (47) to create pCDNA6.2 MR766 clGLuc Intron3127 Pol(-) HDVr.

Huh7 cells were seeded into 6 cm cell culture dishes at a density of 5×10^5^ cells/dish. For siRNA experiments, the cells were transfected with siRNAs 24-hours after seeding of the cells. Twenty-four hours post-transfection, cells were subsequently seeded into 24-well plates at a density of 5×10^4^ cells/well. The following day, 200 ng of the replication-competent and replication-incompetent pCDNA6.2 MR766 clGLuc Intron3127 HDVr, along with appropriate siRNAs (40 nM) or plasmid DNA (500 ng) were transfected using Lipofectamine 3000. At 6, 24, 48- and 96-hours post-transfection, media from wells was collected and stored at −20°C. Assays were performed using NEB BioLux *Gaussia* Luciferase (GLuc) kit (NEB #E3300L) following the stabilized protocol. Briefly, samples and GLuc assay solution were prepared and equilibrated to room temperature. The sample (35 μL) was added to three individual wells in a white 96-well plate. The GLuc assay solution (50 μL) was dispensed into each sample well, shaken for 5 seconds, incubated at room temperature for 30 seconds, and luminescence recorded with an integration time of 10 seconds using a BioTEK Synergy luminometer.

### Statistical Analysis

For RNA quantification of northern blots, ImageQuantTL was used to obtain the mean density of gZIKV, sgZIKV, and actin RNA. RNA levels of ZIKV were initially divided by the mean density of actin and subsequently standardized to the control siGL2 (or 3xFlag-BAP). A two-tailed student T-test comparing control RNA levels to each knockdown (or overexpression) sample was performed using data from three or more biological replicates. Relative viral titers were determined by standardizing the plaque forming units/mL (PFU/mL) for each treatment to the control siRNA (siGL2) or plasmid (p3xFLAG-BAP). A two-tailed student T-test was performed using relative viral titers from three biological replicates. All graphs were generated using Microsoft Excel for Mac 2011. Statistical analysis was performed using StatPlus:mac LE.

## Results

### ZIKV inhibits arsenite-induced SG formation

To determine whether SGs form during ZIKV infection, Huh7 cells were infected with the Cambodian-isolate (160310) at an MOI of 5, and the formation of SGs at 24 hours post-infection was visualized by immunofluorescence and confocal microscopy using an antibody to detect TIA-1, a protein known to facilitate nucleation of SGs (27). ZIKV-infected cells were identified by staining for double-stranded (ds) RNA, a marker of viral replication sites. In mock-infected cells, TIA-1 localized to the nucleus (Figure 1A). Similar to other flavivirus infected cells (30, 31), TIA-1 was mostly localized in the nucleus and we observed ~10% of ZIKV-infected cells containing SG foci in the cytoplasm (Figure 1A and Figure 1C). A lack of SGs during ZIKV-infection suggested ZIKV either inhibited the formation of SGs or promoted the disassembly of SGs. To investigate whether ZIKV inhibited the formation of SGs, we treated cells with 1 mM sodium arsenite for 30 minutes. In mock-infected cells treated with sodium arsenite, we observed 80-100% of the cells containing TIA-1 SGs in the cytoplasm (Figure 1). In contrast, TIA-1-containing SGs were observed in ~24% of ZIKV-infected treated with sodium arsenite (Figure 1B and Figure 1D).

**Figure 1.**
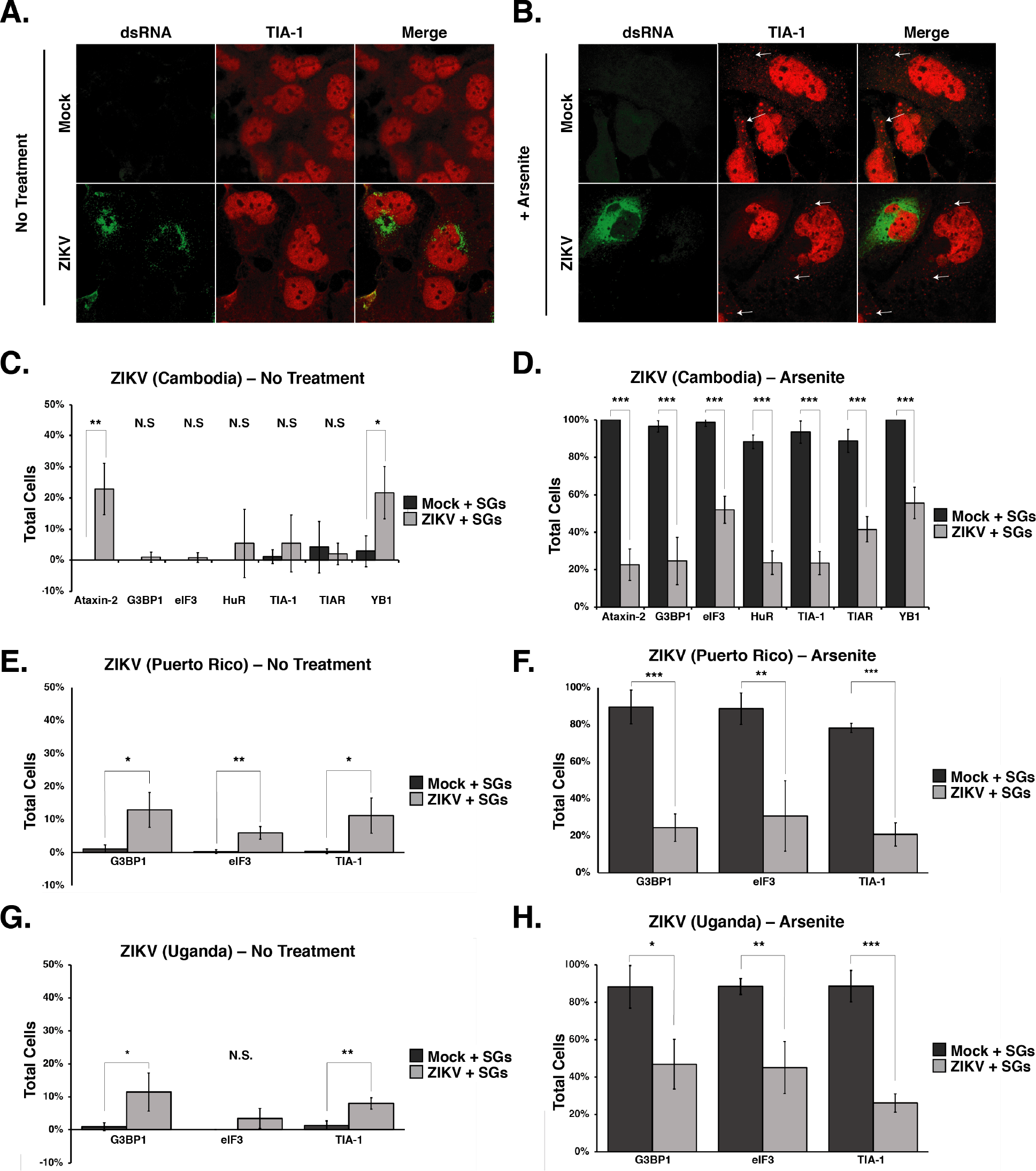
ZIKV restricts the assembly of cellular stress granules. Huh7 cells were either mock-infected or infected with the Cambodia ZIKV-strain (160310) at a MOI of 5. Twenty-four hours post-infection cells were mock-treated or incubated with 1 mM sodium arsenite for 30 minutes, fixed, permeabilized, and analyzed using a 63x oil immersion objective on a Zeiss LSM710 laser scanning confocal microscope. ZIKV-infected cells were detected with mouse-anti-dsRNA antibody (green). The distribution of dsRNA (green) and the stress granule (SG) marker TIA-1 (red) in mock- and ZIKV-infected cells A) not treated and B) treated with sodium arsenite. The white arrows highlight stress granules. Quantification of SGs containing Ataxin-2, G3BP1, eIF3B, HuR, TIA-1, TIAR and YB1 proteins in mock- and ZIKV (Cambodia-160310)-infected which were either C) not treated or D) treated with sodium arsenite. The number of cells containing G3BP1, eIF3, and TIA-1-containing SGs in mock-infected cells or cells infected with a Puerto Rican isolate of ZIKV at a MOI of 5 and either E) not treated or F) treated with sodium arsenite. G-H) SGs in mock-infected cells and cells infected with the original Ugandan isolate of ZIKV at a MOI of 5 were quantified. The number of cells containing G3BP1, eIF3, and TIA-1-SG was quantified from cells that were G) untreated or H) incubated with sodium arsenite. Note, in the original immunofluorescence images, TIA-1 was visualized using an anti-donkey Alexa Fluor® 647 (magenta) anti-goat IgG secondary antibody. In panels A) and B) TIA-1 has been pseudo-colored red. The immunofluorescence images are representative of at least three independent experiments. The errors bars shown on the SG quantification data (Panel 1C, 1D, 1E, 1F, 1G, and 1H) are of the mean ± SD. Significance was determined by two-tailed student T-test (*P<0.05; **P<0.01; ***P<0.001; and N.S. not significant).

We similarly investigated SG formation following infection with the 1947 Uganda-isolate (MR766) and a recent ZIKV strain isolated in Puerto Rico in 2015 (PRVABC59) (44, 45). Similar to Huh7 cells infected with the Cambodia-ZIKV strain (Figure 1C), ~10% of cells infected with MR766 or PRVABC59, had TIA-1-containing SGs (Figure 1E and Figure 1G). Consistent with the number of SGs in cells infected with the Cambodia ZIKV strain (Figure 1D), approximately 20% of cells infected with MR766 or PRVABC formed TIA-1-SGs following treatment with sodium arsenite (Figure 1F and Figure 1H). These data indicate that the three ZIKV strains (Cambodia, Uganda and Puerto Rico) examined restrict the formation of SGs when treated with sodium arsenite and are consistent with SG studies undertaken with WNV, DENV and ZIKV (30, 32, 33, 39, 40).

To determine whether ZIKV inhibited a particular subset of SGs, we next investigated the formation of SGs containing Ataxin-2, G3BP1, eIF3B, HuR, TIAR, and YB1 in mock and ZIKV (Cambodia-160310)-infected cells in the absence or presence of sodium arsenite (Figure 1C and Figure 1D). Quantification of the different SG proteins in mock-infected cells showed that less than 10% of untreated cells contained SGs, while SGs were visible in more than 90% of cells treated with sodium arsenite (Figure 1C and Figure 1D). In contrast, analysis of SGs as visualized using the different SG proteins in cells infected with the Cambodia ZIKV-isolate (160310), showed that 1-25% of untreated cells and 18-56% of sodium arsenite-treated contained SGs (Figure 1C and Figure 1D). We noted that a significant number of ZIKV-infected cells formed Ataxin-2 and YB1 containing SGs in untreated cells, and that the number of cells with SGs containing YB1 increased following sodium arsenite-treatment. Overall, ~20% of ZIKV-infected cells showed sodium arsenite-induced SGs containing common SG components (Figure 1D, Figure 1F and Figure 1H). However, we also observed that 44% of infected cells showed SGs containing the translation initiation factor eIF3B (Figure 1D, Figure 1F and Figure 1H), indicating that the SGs induced in ZIKV-infected cells contain translationally repressed ribonucleoprotein complexes. Together these data show that all three ZIKV strains inhibit the formation of SGs albeit to different extents.

### ZIKV does not dramatically change the abundance or integrity of SG proteins during infection

To determine if the ability of ZIKV to inhibit sodium arsenite-induced SG assembly is a result of promoting proteolytic cleavage of and/or degrading SG-nucleating proteins, we infected Huh7 cells with ZIKV at a MOI of 1 and 5 and investigated the abundance and integrity of select SG proteins at 1, 2, and 3 days post-infection. By western blot analysis we found that the abundance of most SG proteins remained the same between mock- and Cambodia ZIKV-infected samples (Figure 2A), and no cleavage products were identified (Figure 2A and Figure 2B). Interestingly, independent of MOI, the levels of Ataxin-2 increased with ZIKV infection at 1- and 2-days post-infection (Figure 2A). We similarly observed an increase in Ataxin-2 abundance and no change in HuR, TIA-1 and TIAR levels following infections with the Ugandan and Puerto Rican strains (Figure 2B). We also observed a modest decrease in G3BP1 levels (Figure 2A and Figure 2B). While Poliovirus 3C protease has been shown to cleave G3BP1 and block the formation of SGs, we did not observe such a cleavage product (48). It is possible that G3BP1 was cleaved during ZIKV-infection, however the modest decrease in G3BP1 levels likely limited the detection of such a fragment by western blot analysis. Alternatively, G3BP1 might be degraded during ZIKV-infection. Thus, the ability of ZIKV to block the assembly of stress granules was not the result of a change in the abundance or proteolytic cleavage of HuR, TIA-1, and TIAR. However, changes in Ataxin-2 and G3BP1 levels might in part restrict the formation of SGs during ZIKV infection.

**Figure 2.**
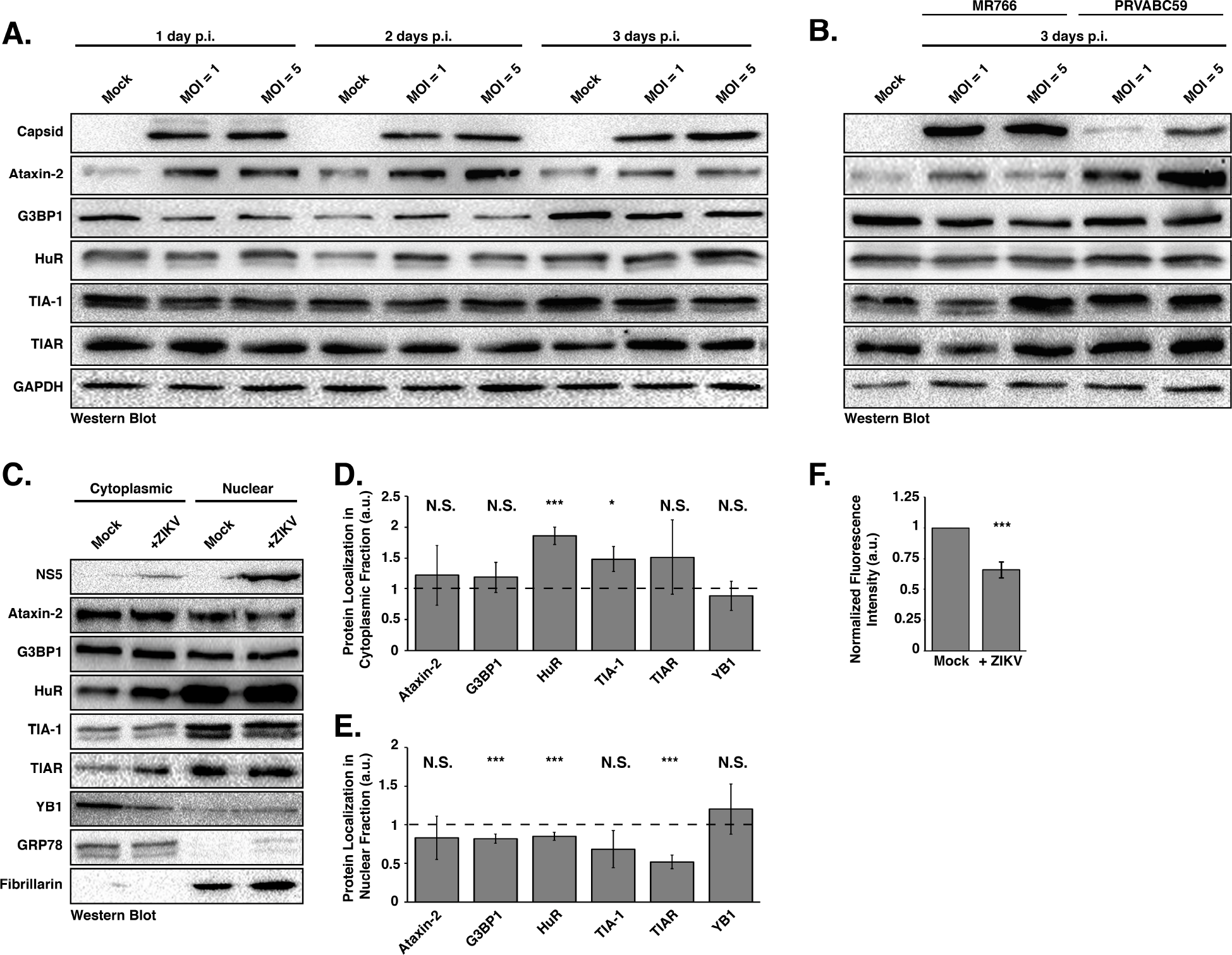
ZIKV infection changes the nuclear and cytoplasmic distribution of stress granule proteins. A) Huh7 cells infected with the Cambodia strain of ZIKV at MOI of 1 or 5 were harvested at 1, 2, and 3 days post-infection. Detection of ZIKV capsid protein by western blot confirmed viral infection. The abundance and integrity of Ataxin-2, G3BP1, HuR, TIA-1 and TIAR proteins was also examined at each time point. The western blot shown is representative of at least three independent experiments. B) Western blot analysis of SG proteins in MR766 and PRVABC59-infected Huh7 cells 3 days post-infection. C) Cytoplasmic and nuclear distribution of Ataxin-2, G3BP1, HuR, TIA-1, TIAR and YB1. Huh7 cells were mock-infected or infected with the Cambodia strain of ZIKV at MOI of 5 and 1-day post-infection the cytoplasmic and nuclear fractions were isolated. The distribution of ZIKV NS5, Ataxin-2, G3BP1, HuR, TIA-1, TIAR, YB1, GRP78 and Fibrillarin was analyzed by western blot. The western blot shown is representative of the three independent experiments performed. Quantification of the distribution of Ataxin-2, G3BP1, HuR, TIA-1, TIAR and YB1 in the D) cytoplasmic and E) nuclear subcellular fractions. The abundance of each SG protein in the cytoplasmic or nuclear fraction was standardized against GRP78 or Fibrillarin, respectively. These values were then normalized against the abundance of the specific SG protein in mock-infected cells, which was assigned the arbitrary unit (a.u.) of one (denoted by dashed line on each graph). The data presented in panels D) and E) were calculated from three independent infections, fractionations and immunoblots. F) Quantification of fluorescence intensity signal of the localization of HuR in the nucleus. Fluorescence intensity was derived in ImageJ using a freehand selection of the nucleus in mock- and ZIKV-infected cells. Values obtained for HuR were divided by the nucleus signal (Hoechst) within mock- and ZIKV-infected cells. Fluorescence intensity of the infected cells were standardized against mock-infected cells and arbitrarily assigned a value of 1. The relative fluorescence intensity signal was calculated from three independent experiments and nine cells were counted per experiment. Two-tailed student T-tests were performed to calculate significance with *P<0.05, ***P<0.001, and N.S. not significant.

### SG proteins are re-localized during ZIKV infection

To investigate whether SG proteins are re-localized during ZIKV infection, we examined the relative distribution of different SG markers in the cytoplasm versus nucleus. In particular, Huh7 cells were either mock- or ZIKV-infected at a MOI of 5, and 1-day post-infection the cytoplasmic and nuclear fractions were isolated and the subcellular distribution of Ataxin-2, G3BP1, HuR, TIA-1, TIAR, and YB1 analyzed by western blot (Figure 2C). To demonstrate effective isolation of the cytoplasmic and nuclear fractions, and to show standardization of the amount of loaded proteins, we examined the localization and abundance of GRP78 (or BiP), an endoplasmic reticulum chaperone protein, and Fibrillarin, the methyltransferase protein that 2’O-methylates ribosomal RNA and is localized in the nucleoli. Western blot detection of ZIKV NS5 confirmed infection, and consistent with immunofluorescence studies (Figure 6A; (49)) showed that ZIKV NS5 was predominantly localized in the nuclear fraction. We semi-quantified the amount of the respective SG protein in each fraction by normalizing the band intensities in the cytoplasmic and nuclear fractions to GRP78 and Fibrillarin, respectively. Normalized values were then standardized to mock samples to obtain a relative change in distribution of each protein. Thus, the ratio of band intensities less-than-one indicate a decrease in the respective fraction, while values greater-than-one represent an enrichment of the SG protein in the subcellular fraction (Figure 2D and Figure 2E). Overall, we did not observe differences in the distribution of Ataxin-2, G3BP1, TIA-1, TIAR and YB-1 between the cytoplasmic and nuclear fractions (Figure 2D and Figure 2E). In contrast however, the relative abundance of HuR in the nuclear fraction decreased, and the concomitant amount of HuR in the cytoplasm increased (Figure 2D and Figure 2E). Indeed, immunofluorescence analysis of HuR localization in mock- and ZIKV-infected cells showed strong nuclear staining, and increased staining in the cytoplasm in ZIKV-infected cells (Figure 5B). We also quantified the fluorescence signal from confocal microscopy images by examining HuR localization in mock- and ZIKV-infected cells. In particular, the fluorescence signal in confocal microscopy images of HuR in the nucleus was quantified. The relative fluorescence intensity signal of HuR, in mock-infected cells was arbitrarily set to 1 (Figure 2F). Using this quantification analysis, a value greater-than-one would indicate increased nuclear localization, and a value less-than-one would indicate decreased localization in the nucleus. Similar to the subcellular fractionation and western blot analysis, we observed a decreased amount of HuR protein in the nucleus in ZIKV-infected cells compared to mock-infected cells (Figure 2E and Figure 2F). These data together show that the subcellular localization of most SG proteins does not significantly change during ZIKV infection. In contrast, however HuR is re-distributed from the nucleus into the cytoplasm in ZIKV-infected cells. That SG proteins, particularly those that promote SG-assembly, do not significantly accumulate in the cytoplasm may in part contribute to the reduction in the number of SGs during ZIKV infection.

### G3BP1 and HuR modulate ZIKV gene expression

To elucidate the role of SG proteins during ZIKV infection we transfected Huh7 cells with siRNAs targeting six different SG proteins and infected with the Cambodian-isolate (160310) at a MOI of 5. Two days post-infection cells were harvested for protein and RNA analysis by western and northern blot, and media was collected for plaque assays. Western blot analysis from at least three independent experiments, showed that the siRNAs effectively depleted the abundance of the targeted SG proteins (Figure 3A). For the majority of the SG proteins examined, siRNA depletion of the individual SG proteins did not impact the levels of other SG markers. We did however observe that the levels of TIA-1 modestly decreased following knockdown of HuR (Figure 3A). TIA-1 levels have similarly been shown to be modulated in cells deficient in another cellular RNA binding protein, namely TIAR (36). In examining the effect of SG protein depletion on ZIKV, the abundance of the ZIKV capsid protein (Figure 3A) did not change following depletion of Ataxin-2. We did however observe a dramatic decrease in viral protein when G3BP1 was depleted, while knockdown of HuR and TIA-1 substantially increased ZIKV capsid protein levels (Figure 3A). Depletion of TIAR and YB1 similarly increased the amount of ZIKV capsid protein, albeit not to the same extent as depletion of HuR and TIA-1 (Figure 3A).

**Figure 3.**
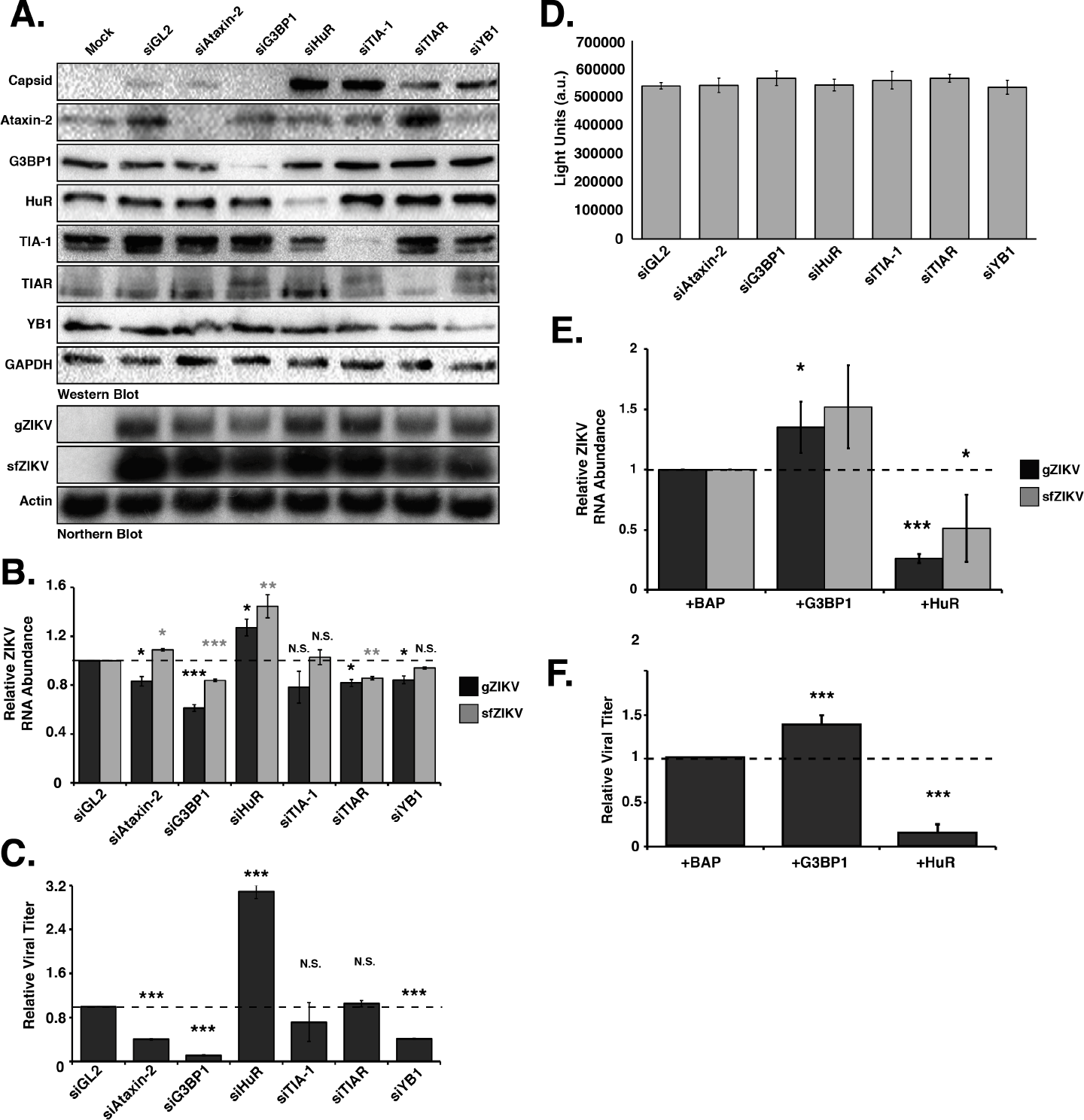
Stress granule proteins promote and limit ZIKV gene expression. Ataxin-2, G3BP1, HuR, TIA-1, TIAR and YB1 proteins were depleted in Huh7 cells using target-specific siRNAs, and then infected 24 hours later at a MOI of 5 with the Cambodia strain of ZIKV (160310). A siRNA targeting *Gaussia* luciferase (siGL2) was used as a control, non-targeting siRNA. Protein, RNA and media were harvested 48 hours post-infection. A) Western blot shows expression of ZIKV capsid, and depletion of each stress granule protein. Expression of GAPDH was used as a loading control, and the western blot shown is representative of at least three independent experiments. The northern blot shows the effect of depleting each stress granule protein on the abundance of the ZIKV genomic (gZIKV) and subgenomic flaviviral (sfZIKV) RNA. Actin mRNA levels were evaluated as a loading control. The northern blot shown is representative of at least three independent experiments. B) Quantification of ZIKV gRNA and sfRNA. ZIKV RNA levels were normalized to actin mRNA, and then represented relative to siGL2 control. C) Viral titers in the extracellular media were determined by plaque assay and relative viral titers were calculated by normalizing titers relative to siGL2 transfected cells. D) A representative assay showing the effect of siRNA depletion of the six SG proteins on cell viability. E) Quantification of ZIKV RNA levels following overexpression of G3BP1-Flag and HuR-Flag. Huh7 cells were transfected with plasmids expressing 3xFlag-Bacterial Alkaline Phosphatase (BAP; control), G3BP1-Flag or HuR-Flag, and then infected with ZIKV (Cambodia-160310) 24 hours post-infection. The effect of overexpressing G3BP1-Flag and HuR-Flag on gZIKV and sfZIKV RNA was evaluated by northern blot. F) Effect of overexpression of G3BP1-Flag and HuR-Flag on viral titers. Data presented in Figure 3 are from at least three independent experiments. Error bars show mean ± SD, and statistical significance was determined using a two-tailed student T-test (*P<0.05; **P<0.01; ***P<0.001; N.S. not significant).

We next examined the abundance of ZIKV genomic RNA (gRNA). By northern blot we observed significant reductions in gZIKV levels following depletion of G3BP1 compared to cells transfected with the control siRNA, siGL2 (Figure 3B). Depletion of Ataxin-2, TIAR and YB1 also decreased gZIKV (Figure 3B). Depletion of TIA-1 did not significantly affect the levels of viral RNA (Figure 3B). Consistent with the increase in ZIKV capsid protein, depletion of HuR significantly increased gZIKV levels (Figure 3B). During flaviviral infections incomplete degradation of the viral gRNA by the cellular 5’-to-3’ exonuclease Xrn1 produces a small noncoding viral RNA or subgenomic flavivirus RNA (sfRNA) that corresponds to the 3’ UTR (50). In our assays we used a northern blot probe to the 3’ UTR and thus also detected the ZIKV sfRNA (Figure 3A and Figure 3B). Overall a decrease in ZIKV gRNA coincided with a decrease in sfRNA, although the abundance of sfRNA was greater than the gRNA, likely because both translating and newly-replicated ZIKV RNA were degraded by Xrn1 (Figure 3B).

We next examined the effect of depleting SG proteins on viral titers by plaque assay (Figure 3C). While, depletion of TIA-1 and TIAR had no effect on the production of infectious virus, depletion of Ataxin-2, G3BP1 and YB1 significantly reduced viral titers. Consistent with the increase in ZIKV protein and RNA following depletion of HuR, we also observed a significant increase in viral titers (Figure 3C).

To ensure the effects on ZIKV following siRNA depletion of the SG proteins were not a result of a change in cell viability, we quantified ATP levels in metabolically active cells from Huh7 cells transfected with the respective siRNAs. The cell viability assay showed no significant differences in the metabolic activity in cells depleted of Ataxin-2, G3BP1, HuR, TIA-1, TIAR and YB1, indicating that the changes in ZIKV protein, RNA, and viral titers were the consequence of knockdown of the specific SG protein (Figure 3D).

Because siRNA-depletion of G3BP1 and HuR showed the most dramatic effect on ZIKV gene expression, we chose to focus on these two SG proteins in subsequent experiments. To confirm the effect of G3BP1 and HuR on ZIKV gene expression, we overexpressed G3BP1-Flag and HuR-Flag in ZIKV-infected cells and examined viral RNA levels and viral titers (Figure 3E and Figure 3F). Analysis of viral RNA abundance showed that overexpression of G3BP1 and HuR significantly increased and decreased ZIKV RNA, respectively (Figure 3E). Consistent with the effects on ZIKV RNA, we observed a corresponding increase and decrease in viral titers following overexpression of G3BP1 and HuR (Figure 3F). Together these data indicate TIA-1, TIAR and YB1 regulate translation of the ZIKV polyprotein, and have a role in replication of the viral genome (Figure 3A and Figure 3B). The significant decrease in viral titer following depletion of YB1 further suggest a role for this cellular RNA-binding protein in virus assembly (Figure 3C). Ataxin-2 and G3BP1 are proviral cellular factors as depletion of Ataxin-2 and G3BP1 decreased ZIKV RNA levels (Figure 3A and 3B), and overexpression of G3BP1 increased viral RNA (Figure 3E), which in turn affected the production of infectious particles (Figure 3F). Finally, HuR exhibits antiviral activity as depletion of this RNA-binding protein significantly increased ZIKV protein and RNA levels and viral titers, and overexpression of HuR-Flag showed a reciprocal effect on ZIKV gene expression (Figure 3).

### Changing the abundance of G3BP1 and HuR affects ZIKV replication

Depletion of G3BP1 and HuR significantly decreased and increased ZIKV gene expression respectively (Figure 3), suggesting a role for these SG proteins in ZIKV translation and/or replication. To decipher the function of G3BP1 and HuR in ZIKV gene expression, we used the replication-competent (WT) and -deficient (Pol-) *Gaussia* luciferase (GLuc) ZIKV reporter plasmid (pCDNA6.2 MR766 clGLuc Intron3127 HDVr, Figure 4A). These pCDNA6.2 MR766 clGLuc Intron3127 HDVr constructs were derived from the cDNA clone containing the full-length genome of the Uganda MR766 strain under the control of the cytomegalovirus (CMV) promoter (Figure 4A) (47). Specifically, the full-length GLuc reporter gene, along with its signal sequence and the sequence for the foot-and-mouth disease virus (FMDV) 2A peptide, was cloned as a translational fusion following the first 20 amino acids of the capsid protein, and the full-length viral polyprotein was cloned downstream of GLuc. To avoid aberrant genome cyclization during replication dues to duplicated 5’ cyclization elements, the flavivirus cyclization sequences in full length capsid coding region were disrupted by silent mutagenesis. Following transfection of this reporter construct, transcription and 5’-end capping of ZIKV genomic RNA are directed by the host, and the hepatitis D virus ribozyme (HDVr) at the end of the 3’ UTR cleaves the ZIKV genomic RNA to create an authentic 3’ end (Figure 4A) (47). Using GLuc expression as a proxy for viral RNA, translation of capped ZIKV RNA is detected 6 hours post-transfection. Translation of the ZIKV polyprotein, including the synthesis of NS5, the RNA-dependent-RNA polymerase (RdRp), directs replication of the reporter genome such that the increase in viral RNA as a result of replication is observed at 48- and 96-hours post-transfection. We also expressed a ZIKV GLuc reporter genome that contained a mutation in the RdRp active site (GDD-to-GNN; pCDNA6.2 MR766 clGLuc Intron3127 Pol(-) HDVr) which rendered this mutant GLuc reporter genome replication-deficient (Pol-). Expression of the Pol(-) reporter genome showed a clear difference between replicating WT genomes and ZIKV RNA transcribed from the transfected plasmid at 48- and 96-hours post-transfection (Figures 4B-4E).

**Figure 4.**
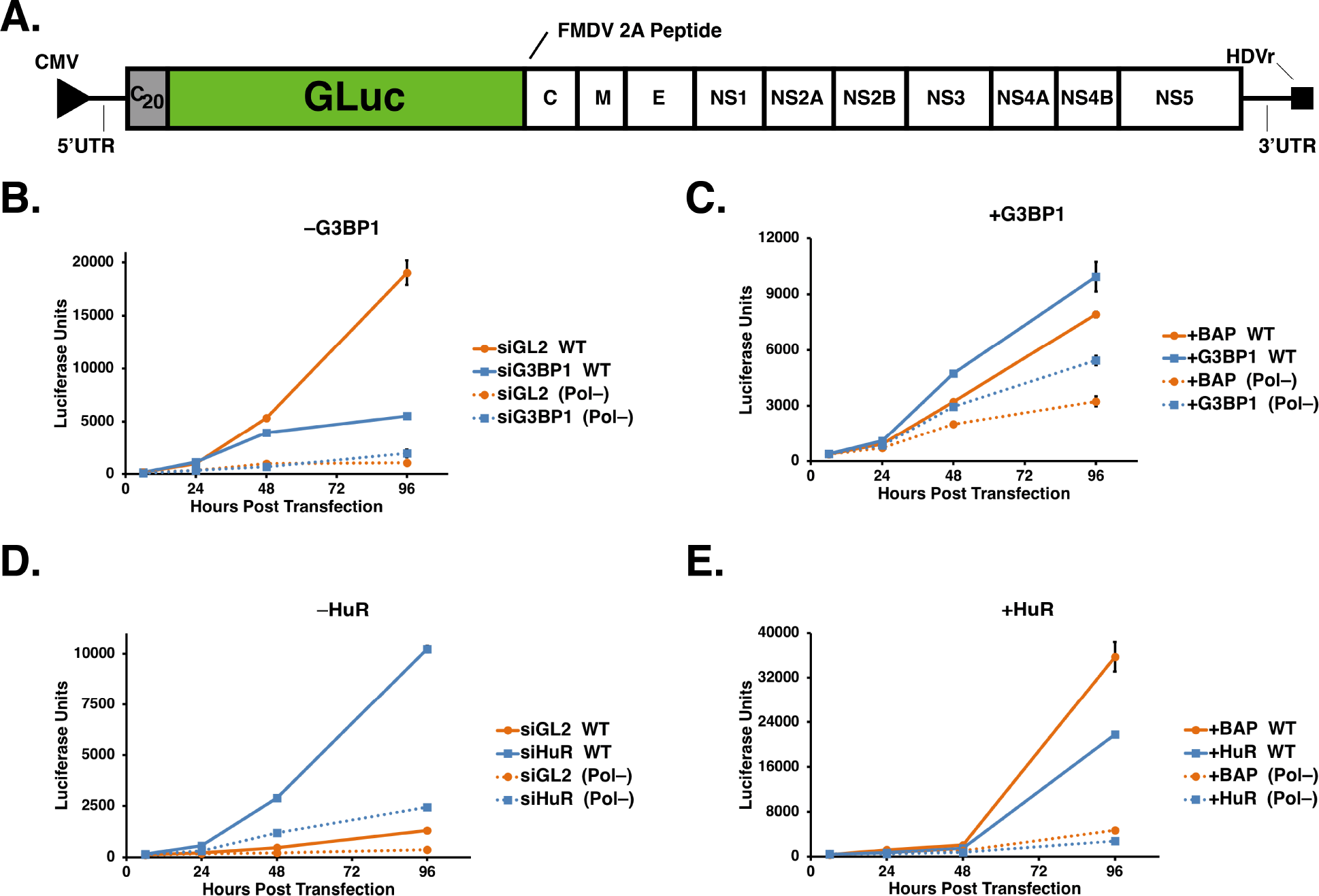
G3BP1 and HuR modulate ZIKV replication. A) Schematic of the ZIKV MR766 *Gaussia* luciferase reporter genome within the pCDNA6.2 MR766 clGLuc Intron3127 HDVr plasmid showing the 5’ and 3’ UTRs, the mature viral proteins within the single-open reading frame, as well as the position of the *Gaussia* luciferase gene within the MR766 genome. Two elements within the plasmid namely the CMV promoter and the HDVr, which creates an authentic 3’ UTR in the genome, are also denoted. B & D) Huh7 cells were first transfected with either the control or target-specific siRNAs, and then transfected with the same siRNAs and the pCDNA6.2 MR766 clGLuc Intron3127 HDVr WT replication-competent or Pol(-) replication-deficient plasmids. GLuc activity in the media of transfected cells was assayed at 6, 24, 48, and 96 hours post-transfection. C & E) Huh7 cells were transfected with p3xFlag-BAP, pG3BP1-Flag, pHuR-Flag and WT and Pol(-) pCDNA6.2 MR766 clGLuc Intron3127 HDVr. At 6, 24, 48, and 96 hours post-transfection media from the transfected cells was collected and GLuc activity measured. B) Effect of G3BP1 knockdown on ZIKV-GLuc genome expression. C) Effect on ZIKV-GLuc reporter genome expression following overexpression of 3xFlag-BAP (control) and G3BP1-Flag. D) Effect of depleting HuR on ZIKV-GLuc genome expression. E) Effect of overexpressing 3xFlag-BAP and HuR-Flag on ZIKV-GLuc genome expression. The data shown in Figure 4B-4E are from a single experiment and are representative of at least three independent experiments. Error bars indicate mean ± SD.

To elucidate effects on translation and replication we examined expression of the ZIKV GLuc reporter genome following siRNA depletion or overexpression of G3BP1 and HuR (Figures 4B-4E). RNAi depletion or overexpression of G3BP1 and HuR did not significantly affect GLuc expression at 6- and 24-hours post-transfection of both the ZIKV WT and Pol(-) reporter genomes (Figures 4B-4E), indicating that these two SG proteins do not affect translation of the ZIKV genome (Figures 4B-4E). At 96 hours post-transfection of the WT reporter genome however, siRNA depletion and overexpression of G3BP1 notably decreased and increased GLuc expression respectively, showing that G3BP1 facilitates ZIKV replication (Figure 4B and Figure 4C). Curiously, overexpression of G3BP1 also increased GLuc expression from the ZIKV Pol(-) reporter genome (Figure 4C). Since this ZIKV Pol(-) reporter genome is replication deficient, the increased levels of host-transcribed ZIKV mRNA might be the result of G3BP1 binding (39) and stabilizing the viral mRNA. In contrast to the effect of G3BP1, at 96 hours post-transfection depletion of HuR dramatically increased GLuc expression (Figure 4D). Reciprocally, overexpression of HuR decreased luciferase units compared to expression of the 3xFlag-BAP control (Figure 4E), indicating that HuR negatively impacts the ZIKV replication. Taken together these data support a role for G3BP1 and HuR in replication of the ZIKV genome.

### G3BP1 and HuR localize with ZIKV replication complexes

To further demonstrate a role for G3BP1 and HuR in ZIKV replication, we next examined the localization of these two SG proteins at replication complexes (Figure 5). We first examined the localization of G3BP1 and HuR at replication complexes by immunofluorescence analysis. In particular, we used antibodies to dsRNA and ZIKV NS5 protein as markers for replication sites (Figure 5A and Figure 5B). While dsRNA is a replication intermediate of negative-and positive-sense dsRNA, NS5 is critical for the replication of the ZIKV RNA as it functions as the RdRp and the methyltransferase that caps the 5’ end of the ZIKV genome (1). G3BP1 shows diffuse cytoplasmic staining in mock-infected cells, however in ZIKV-infected cells we observed strong co-localization of G3BP1 with dsRNA (Figure 5A), and some co-localization with NS5 in the cytoplasm (Figure 5A). In mock-infected cells, HuR was predominantly localized in the nucleus (Figure 5B). Consistent with the subcellular fractionation data (Figure 2B and Figure 2C) we observed an increase in the localization of HuR in the cytoplasm of ZIKV-infected cells (Figure 5B). Although HuR did not co-localize with dsRNA in ZIKV-infected cells, we did note HuR staining adjacent to ZIKV replication sites (Figure 5B). HuR showed similar localization with NS5 in the cytoplasm (Figure 5B). The localization of G3BP1 and HuR with respect to ZIKV replication sites support a role in the synthesis of viral RNA.

**Figure 5.**
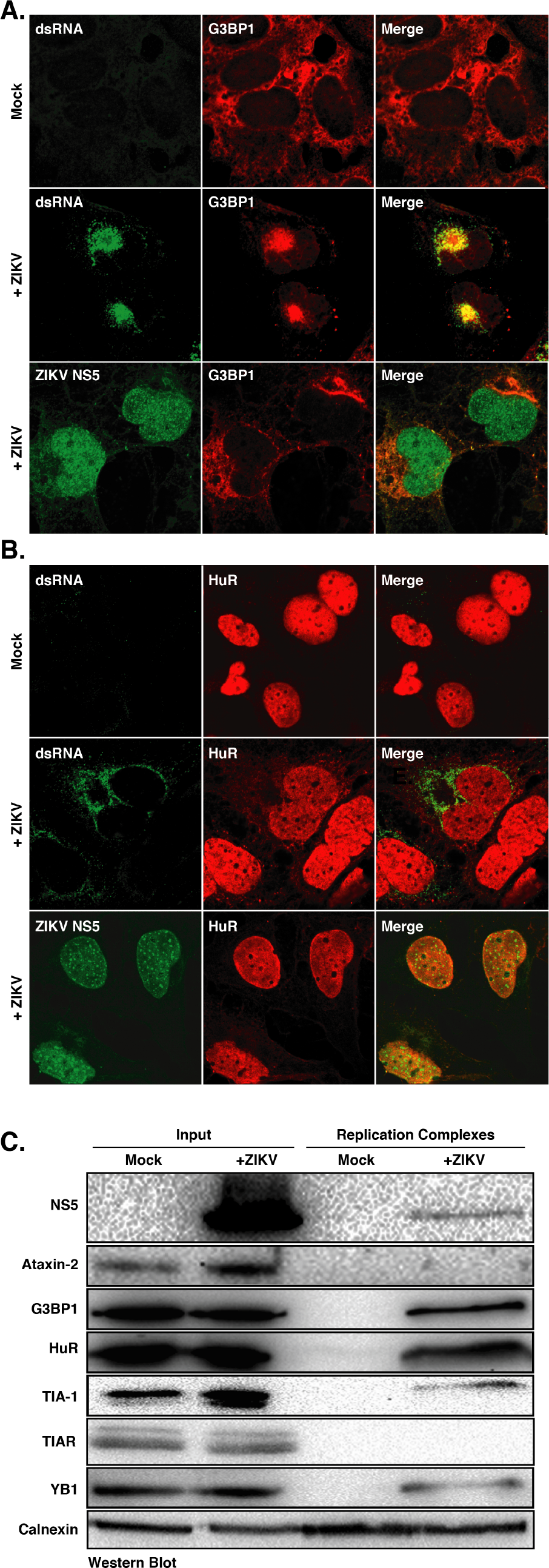
G3BP1 and HuR localize with ZIKV replication complexes. A-B) Huh7 cells were either mock-infected or infected with the Cambodia ZIKV strain at a MOI of 5. One day post-infection cells were fixed, permeabilized, and prepared for confocal imaging. Immunofluorescence images are representative of at least three independent experiments. A) Immunofluorescence image showing G3BP1 (red) localization in mock-infected cells and with viral replication sites as visualized by the staining with antibodies to dsRNA and NS5 (green) in ZIKV-infected cells. B) Localization of HuR (red) in mock-infected cells and with dsRNA and NS5 (green) in ZIKV-infected cells. C) Western blot analysis of ZIKV NS5, Ataxin-2, G3BP1, HuR, TIA-1, TIAR, YB1, and calnexin associated with replication complexes isolated from mock- and ZIKV-infected Huh7 cells. The western blot is representative of three independent experiments.

**Figure 6.**
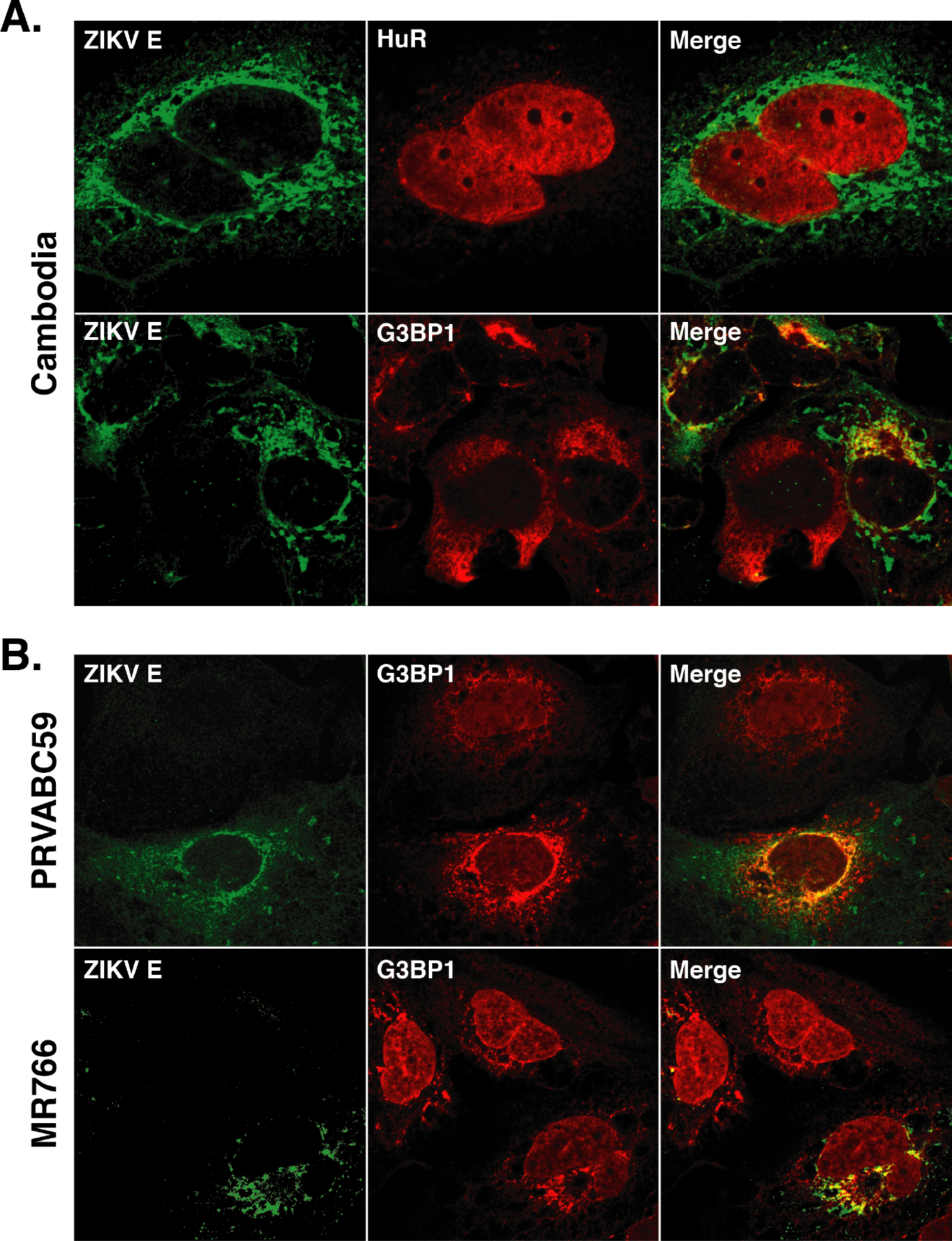
G3BP1 co-localizes with ZIKV E protein. One day post-infection Huh7 cells infected with ZIKV at a MOI of 5 were fixed, permeabilized, and prepared for confocal imaging. A) Localization of ZIKV E protein with HuR and G3BP1 following infection with the Cambodia strain of ZIKV. B) Localization of ZIKV E protein with G3BP1 following infection with the Puerto Rican (PRVABC59) and Ugandan (MR766) strains of ZIKV, respectively. The immunofluorescence images shown are representative of at least three independent experiments.

Similar to other immunofluorescence studies (49), we observed strong nuclear and weak cytoplasmic localization for the ZIKV NS5 protein (Figure 5A and Figure 5B). Such localization was consistent with the cytoplasmic/nuclear fractionation data (Figure 2B, Figure 2C and Figure 2D). Because both NS5 and HuR are largely nuclear, we anticipated that these proteins might co-localize. Although immunofluorescence analysis showed diffuse nuclear staining for both NS5 and HuR, HuR did not co-localize with NS5 foci in the nucleus (Figure 5B). Both HuR and flavivirus NS5 proteins have been shown to affect RNA splicing (51). While it is possible that HuR and NS5 together in the nucleus might modulate processing of cellular RNAs, further interaction studies would need to be undertaken to verify such a role.

To corroborate the immunofluorescence analyses and role for G3BP1 and HuR in ZIKV replication, we also isolated replication complexes from mock- and ZIKV-infected Huh7 cells by sucrose gradient ultracentrifugation, and then investigated the localization of select SG proteins with replication complexes. Western blot analysis showed the isolated fractions were derived from the endoplasmic reticulum as shown by calnexin immunostaining in both mock and ZIKV-infected fractions, and NS5 protein was only isolated in membrane fractions isolated from ZIKV-infected cells. G3BP1 and HuR were isolated with the replication complexes (Figure 5C). TIA-1, TIAR and YB1 have previously been shown to affect WNV replication (30, 36), we only observed a small amount of TIA-1 and YB1 in the enriched membranes (Figure 5C). While we noted an increase in the Ataxin-2 levels in ZIKV infected cells (Figure 2A), Ataxin-2 did not sediment with replication complexes (Figure 5C). That G3BP1 and HuR were isolated with replication complexes (Figure 5C) further indicates that G3BP1 and HuR function to modulate ZIKV replication (Figure 4).

### G3BP1 co-localizes with ZIKV envelope protein

Because ZIKV replication and assembly occur in close proximity on cellular membranes, and G3BP1 was previously shown to interact with the capsid protein (39), we also investigated whether G3BP1 and HuR co-localized with ZIKV assembly sites. In particular, we examined the co-localization of both proteins with ZIKV envelope (E) protein by immunofluorescence analysis (Figure 6). Similar to the localization of HuR with dsRNA and ZIKV NS5 protein (Figure 5B) a small amount of HuR co-localized with ZIKV envelope protein (Figure 6A). Interestingly, G3BP1 showed co-localization with the envelope protein in cells infected with the Cambodia ZIKV strain (Figure 6A) and with the Puerto Rican (PRVABC59) and Ugandan (MR766) strains of ZIKV (Figure 6B). The co-localization of G3BP1 with the ZIKV envelope protein together with the siRNA studies showing that depletion of G3BP1 modestly decreased ZIKV RNA by ~40% (Figure 3B), while viral titers were dramatically reduced (Figure 3C), raise the possibility that G3BP1 might function more broadly than previously recognized. Specifically, G3BP1 promotes replication of the ZIKV genome and maybe also the assembly of new viral particles.

## Discussion

The formation and sequestration of components of the translation machinery in SGs function as an intrinsic antiviral mechanism to restrict viral translation and/or replication. Flaviviruses have been shown to limit stress granule assembly by affecting cellular pathways. Specifically WNV was shown to modulate the cellular redox response by increasing the levels of the transcription factors that activate antioxidant genes and increase reduced glutathione levels (32). DENV modulates stress granule assembly by changing the phosphorylation of the cap-binding protein eIF4E by altering p38-Mnk1 signaling to affect cellular translation (33). More recently, ZIKV was shown to increase the rate of eIF2α dephosphorylation which would promote translation and the rapid disassembly of stress granules (40). Interestingly, Hou *et al.*, also showed that specific viral proteins impact cellular translation and the formation of stress granules (39). We similarly observed fewer SGs in ZIKV-infected cells, and that the assembly of SGs following sodium arsenite-treatment was inhibited during ZIKV infection (Figure 1). Proceeding depletion of TIAR, G3BP1 and Caprin-1, Hou *et al.*, also found decreased ZIKV viral titers (39). In our study we present a systematic investigation of the role of six different SG proteins on ZIKV protein and RNA abundance, production of infectious particles, the co-localization with ZIKV and effects on replication. Although G3BP1 from A549-infected cells co-immunopreciptated ZIKV RNA (39), we specifically show that G3BP1 is in replication complexes (Figure 5) and promotes synthesis of ZIKV RNA (Figure 4B and Figure 4C). Hou *et al.*, also demonstrated that G3BP1 interacted with overexpressed Flag-tagged ZIKV capsid protein (39). We found that in ZIKV-infected cells G3BP1 co-localized with ZIKV envelope protein which raises the possibility that G3BP1 may facilitate virion assembly in addition to its role in ZIKV replication (Figure 6). Excitingly, we show that HuR functions to limit ZIKV replication, which to our knowledge has not previously been shown for this cellular RNA binding protein. Our systematic analysis of SG proteins during ZIKV infection is significant particularly as G3BP1, HuR and other SG proteins have an intimate role in neurodevelopment and function. The modulation of SGs and SG proteins by ZIKV may have a more significant impact on normal gene regulation that contributes to ZIKV pathogenesis.

Ataxin-2 is a cytosolic SG protein that is associated with the neurodegenerative disorder spinocerebellar ataxia type-2 (SCA2) (52, 53). During ZIKV infection we observed an increase in the abundance of Ataxin-2 (Figure 2A). Furthermore, RNAi studies showed that depletion of Ataxin-2 did not affect ZIKV capsid levels (Figure 3A) but did decrease the amount of viral RNA and viral titers (Figure 3B and Figure 3C). Ataxin-2 levels have been shown to affect the abundance of the poly(A) binding protein (PABP) (53), where elevated levels of Ataxin-2 decreased the amount of PABP (53). Such changes in PABP would be expected to negatively affect translation of cellular mRNAs and promote ZIKV gene expression. Moreover, Ataxin-2 interacts with the cellular RNA DEAD-box helicase DDX6 (53), which we have previously shown to be required for ZIKV gene expression (54). Therefore, the impact on ZIKV RNA and production of infectious particles (Figure 3B and Figure 3C) following RNAi depletion of Ataxin-2 might also be the result of changes to DDX6. Analysis of SCA2 brains showed an increase in Ataxin-2 levels and that Ataxin-2 localized in intranuclear inclusions in 1-2% of neurons (55, 56). Our data showing that Ataxin-2 modulates ZIKV gene expression in the hepatocellular carcinoma cell line Huh7 is exciting and additional studies in neuronal cells would illuminate the role of this protein in cellular dysfunction and neurodegeneration following intrauterine ZIKV infection.

The RNA-binding proteins TIA-1 and TIAR are broadly expressed in cells and tissues (57). These proteins are predominantly localized in the nucleus (Figure 1, and data not shown), have been shown to modulate translation of specific cellular mRNAs (58) and are critical nucleators of stress granules in the cytoplasm (27). During WNV and DENV infection TIA-1 and TIAR co-localize at replication sites, and bind to the 3’ stem loop region in the negative-strand of WNV RNA to promote replication (30, 36, 59). We determined by western blot analysis that TIA-1, but not TIAR, localized in isolated replication complexes (Figure 5C). Because TIA-1 and TIAR were previously shown to promote WNV replication we also investigated the role of TIA-1 and TIAR during ZIKV infection. Similar to Hou *et al.* (39), we found that decreasing the levels of TIA-1 had no effect on viral titers (Figure 3C). In contrast to Hou *et al.* (39), we found that that depletion of TIAR did not affect ZIKV titers (Figure 3C). Our RNAi studies also demonstrated that following the reduction of TIA-1 and TIAR levels, the abundance of ZIKV capsid protein increased (Figure 3A), and depletion of TIAR decreased ZIKV RNA levels (Figure 3A and Figure 3B), suggesting that, similar to tick-borne encephalitis virus, TIA-1 and TIAR are re-localized to perinuclear regions of replication to modulate translation of the ZIKV polyprotein (60). While the role of TIA-1 and TIAR in our studies in Huh7 differs from the study by Hou and colleagues which were undertaken in A549 cells (39), the impact of TIA-1 and TIAR might be better elucidated in neuronal cells where a link between TIA-1, the microtubule protein Tau and the neurodegenerative tauopathies has been described (61, 62).

Y box-binding protein-1 (YB1) modulates translation and stabilizes cellular mRNAs (58) and also localizes in processing bodies and stress granules (63). YB1 has been shown to bind the DENV 3’ UTR to decrease DENV translation (64). In our study, siRNA depletion of YB1 increased the levels of ZIKV capsid protein (Figure 3A) indicating that YB1, similar to the role in DENV infection, negatively regulates ZIKV translation. We also observed that YB1 is localized in replication complexes (Figure 5C) and that depleting YB1 levels decreases ZIKV RNA levels (Figure 3B). Notably however, the modest effects of YB1 reduction on ZIKV protein and RNA levels differ from the significantly decreased viral titers (Figure 3C). YB1 was previously shown to interact with the hepatitis C virus NS3/4A protein, and other cellular factors, to modulate the balance between replication and the assembly of HCV infectious particles (65, 66), and additional studies with YB1 may reveal a similar function during ZIKV infection.

G3BP1, similar to TIA-1, is an important nucleator of SGs (26, 28, 67). Interestingly, the role of G3BP1 varies in viral infections from limiting alphavirus replication (68–70) to interacting with the HCV RdRp NS5B and enhancing the production of HCV infectious virus particles (43, 71, 72). In DENV-infected cells, G3BP1 interacts with the sfRNA (37, 73), and sequestration of G3BP1 by the sfRNA was shown to inhibit translation of interferon-stimulated genes (38). Hou *et al.*, showed that siRNA depletion of G3BP1 decreased ZIKV titers and that G3BP1 also bound ZIKV RNA (39). In addition to the dramatic reduction in ZIKV titers following depletion of G3BP1 (Figure 3C) we also observed a decrease ZIKV protein and RNA levels (Figure 3A and Figure 3B). Together these data suggested a role for G3BP1 in ZIKV translation, replication, and/or assembly. In our study we also used the ZIKV MR766 GLuc reporter genome (Figure 4B and Figure 4C) and showed a specific role in ZIKV replication, which was further supported by the localization of G3BP1 with ZIKV replication complexes (Figure 5C) and co-localized with dsRNA (Figure 5A). We also observed that ZIKV envelope protein (Figure 6) co-localized with G3BP1 suggesting that in addition to facilitating replication of the ZIKV genome, G3BP1 may have a role in assembly of new virus particles. Hou and colleagues report that G3BP1 binds ZIKV gRNA and that G3BP1 and Caprin-1 interact with the ZIKV capsid protein to increase ZIKV gene expression. While further mechanistic studies are required to demonstrate a role for G3BP1 in ZIKV assembly, it is possible that a multi-protein complex between ZIKV RNA, capsid protein, G3BP1 and Caprin-1 may shift the equilibrium from replication of the viral genome towards localization with the ZIKV envelope protein and the assembly of infectious ZIKV particles.

Human antigen R (HuR) is a member of the ELAVL RNA-binding protein family and is ubiquitously expressed (74). Notably, the three additional members of the ELAVL family (HuB/C/D) are highly expressed in the brain (74). HuR binding to poly-U and AU-rich elements within the 3’ UTR increases mRNA stability (75, 76). During infection with the single-stranded positive-sense alphaviruses Sindbis virus (SINV) and Chikungunya virus (CHIKV), HuR was dramatically re-localized from the nucleus into the cytoplasm (77, 78). By subcellular fractionation and western blot analysis we similarly observed that HuR is re-distributed into the cytoplasm (Figure 2B), albeit not to the same extent as during alphavirus infection (77, 78). HuR is recruited to alphavirus RNA via the U-rich elements within the 3’ UTR and this interaction stabilizes and protects the viral RNA from the cellular mRNA decay machinery (78). Although HuR has been shown to promote alphavirus mRNA stability (77, 78) and HCV IRES-mediated translation (43, 79) our study indicates an antiviral role for HuR during ZIKV infection (Figure 3), which to our knowledge has not previously been shown. In particular we found that HuR modulates ZIKV replication (Figure 4D and Figure 4E) which resulted in increased ZIKV protein and RNA levels and viral titers (Figure 3). The role of HuR in ZIKV replication was further supported by the detection of HuR in isolated replication complexes (Figure 5C). Curiously, immunofluorescence studies showed that HuR in the cytoplasm was localized adjacent to dsRNA and ZIKV NS5 protein staining (Figure 5B). We note that the localization of HuR in replication complexes when analyzed in isolated replication complexes versus immunofluorescence analysis do not strongly correlate. However, it is important to highlight that HuR is strongly localized in the nucleus (Figure 5B and 6A) and this nuclear staining likely overshadows the re-localization of HuR into the cytoplasm and replication complexes. Indeed, immunofluorescence analysis of ZIKV NS5 protein similarly showed strong nuclear and weak cytoplasmic localization (Figure 5B), despite being the RdRp responsible for replicating the ZIKV RNA genome. A role for HuR in ZIKV replication was further supported by siRNA depletion of HuR and overexpression of HuR-Flag that increased and decreased replication of the MR766 GLuc reporter expression respectively (Figure 4D and Figure 4E). The mode by which HuR limits ZIKV replication remains to be elucidated. A bioinformatic analysis of the ZIKV 3’ UTRs identified putative HuR binding sites (data not shown), raising the possibility that similar to SINV (78), HuR interacts with ZIKV RNA to alter the subcellular localization of HuR, and directly affect ZIKV replication, possibly by destabilizing, rather than stabilizing, the viral RNA. Alternatively, the consequence on ZIKV replication might be indirect where HuR re-localization affects cellular RNA homeostasis or ribostasis. Indeed, re-localization of HuR by SINV into the cytoplasm was shown to decrease cellular mRNA stability and alter mRNA splicing and nuclear polyadenylation (80). Interestingly, another cell RNA-binding protein Fragile X mental retardation factor (FMRP) was shown to bind ZIKV sfRNA and limit viral infection (81). It will be interesting to determine the mechanism by which HuR functions during ZIKV, and whether this regulatory mode is similar to FMRP. Such studies will provide new insights into viral-host interactions, particularly for RNA-binding proteins that function as restriction factors.

Beyond aggregating stalled translation complexes, RNA binding proteins in stress granules also localize in neuronal granules. These granules regulate neuronal growth and synaptic plasticity (82, 83). Interestingly, a number of stress granule proteins such as Ataxin-2, G3BP1, TIA-1 and HuR are known to contribute to different neuropathologies (41, 83). While transcriptional changes have been reported to contribute to the neurological and developmental defects observed following intrauterine ZIKV-infection (84–87), the role of stress granules and granule proteins should not be overlooked. By limiting the formation of stress granules, and re-localizing and subverting specific stress granule proteins, alterations in RNA homeostasis, such as changes in RNA splicing, RNA stability and translation, could also contribute to ZIKV neuropathologies. To this end, elucidating the interactions of stress granule proteins and the cellular consequences particularly in neuronal cells could provide new insights into the pathomechanisms underlying ZIKV congenital disease.

## Acknowledgements

We thank Dr. Brett Lindenbach (Yale School of Medicine) for ZIKV Cambodian-isolate (160310 and Ugandan-isolate (MR766), and Dr. Laura Kramer (Wadsworth Center, NYSDOH) and the CDC for the Puerto Rican-isolate (PRVABC59). We also thank members of the Pager lab, Marlene Belfort, Gabriele Fuchs and Ing-Nang Wang for valuable comments and suggestions on the manuscript.

